# A circadian output circuit controls sleep-wake arousal threshold in *Drosophila*

**DOI:** 10.1101/298067

**Authors:** Fang Guo, Meghana Holla, Madelen M. Díaz, Michael Rosbash

## Abstract

The *Drosophila* core circadian circuit contains distinct groups of interacting neurons that give rise to diurnal sleep-wake patterns. Previous work showed that a subset of Dorsal Neurons 1 (DN1s) are sleep-promoting through their inhibition of activity-promoting circadian pacemakers. Here we show that these anterior-projecting DNs (APDNs) also “exit” the circadian circuitry and communicate with the homeostatic sleep center in higher brain regions to regulate sleep and sleep-wake arousal threshold. These APDNs connect to a small discrete subset of tubercular-bulbar neurons, which are connected in turn to specific sleep-centric Ellipsoid Body (EB)-Ring neurons of the central complex. Remarkably, activation of the APDNs produces sleep-like oscillations in the EB and also raises the arousal threshold, which requires neurotransmission throughout the circuit. The data indicate that this APDN-TuBu_sup_-EB circuit temporally regulates sleep-wake arousal threshold in addition to the previously defined role of the TuBu-EB circuit in vision, navigation and attention.

## Introduction

Sleep is a common biological state observed in many if not most animals, in which they exhibit periods of prolonged immobility, reduced responsiveness to the environmental input, and certain patterns of central brain activity (Allada and Siegel, 2008; Campbell and Tobler, 1984; Greenspan et al., 2001). The function of sleep is still uncertain and has been a topic of scientific inquiry in mammals for decades. More recently these functions have also been studied in *Drosophila melanogaster*, a model organism for human function because of its highly conserved genes and relatively simple nervous system (Allada and Siegel, 2008; Celniker and Rubin, 2003; Hendricks et al., 2000; Rubin et al., 2000; Shaw et al., 2000)

Sleep is highly regulated by two independent systems: a homeostatic mechanism and the circadian clock (Borbely et al., 2016; Ho and Sehgal, 2005). The homeostatic mechanism determines the duration of sleep based on prior sleep-wake history (Borbely and Achermann, 1999). The endogenous circadian clock in contrast ignores past history and maintains a regular 24-hour sleep/wake cycle, effectively directing the time in which organisms prefer to sleep. The circadian clock regulates behavior and has three main components: the core clock, which keeps time and coordinates the clock circuit; input pathways, which adapts and synchronizes the core clock to the environment changes; and output pathways, which transmit information to higher brain regions to control behavior at certain circadian times (Allada and Chung, 2010).

The core clock brain network is composed of 75 pairs of neurons symmetrically located in both hemispheres, which express essential circadian genes such as *period* and *timeless* to generate 24-hour rhythms(Dubowy and Sehgal, 2017). Clock neurons can be categorized as the small and large ventral lateral neurons (LNvs), the dorsal lateral neurons (LNds), dorsal neurons (DN1s, DN2s, DN3s), and the lateral posterior neurons (LPNs) (Allada and Chung, 2010; Nitabach and Taghert, 2008). The small LNvs (s-LNvs) are morning cells (M-cells) that regulate *Drosophila* morning locomotor activity and free-running rhythms, whereas the fifth s-LNv and the LNds are evening cells (E-cells) that regulate evening activity (Grima et al., 2004; Helfrich-Forster, 2000). A subset of glutamatergic DN1s have an additional function (Zhang et al., 2010a; Zhang et al., 2010b), which is to negatively feedback onto the M and E cells to promote sleep, especially during midday (Guo et al., 2016). This feedback and the clock neuronal circuit more generally maintain the normal *Drosophila* sleep/wake cycle.

Compared to the detailed understanding of the fly circadian circuit in sleep regulation, the neural and molecular underpinnings of sleep outside of this circuit are less well defined. Several widely dispersed brain regions have been proposed as centers for sleep/wake regulation, which include the mushroom body (Guo et al., 2011; Joiner et al., 2006; Pitman et al., 2006), the dorsal fan-shaped body (dFSB) (Donlea et al., 2011; Liu et al., 2012), and the ellipsoid body (EB) (Donlea et al., 2018; Liu et al., 2016). In addition, a ring substructure of the EB, the EB-R2 neurons, appears to play an essential role in regulating sleep (Liu et al., 2016).

Importantly, there has been no comprehensive functional or anatomical investigation of a connection between the *Drosophila* circadian circuit and these other sleep-relevant fly brain regions. The circadian circuit is anatomically dispersed with most of the important circadian neurons on the lateral or dorsal surface of brain; they are therefore well-positioned to sense the environment, for example light and temperature fluctuations (Yadlapalli et al., 2018; Zhang et al., 2010b). However, it is still unclear how this sensory information is integrated with circadian timekeeping and then processed to change central brain activity patterns and most importantly to influence the status of wake/sleep and arousal levels.

In this paper, we show that circadian sleep and wake signals from the DN1 neurons are encoded and conveyed by two separate circadian output circuits with distinct projection patterns. The wake-promoting pathway projects to the dorso-medial protocerebrum, the so-called pars intercerebralis (PI), whereas the sleep-promoting DNs send neurites to the anterior region and make synapses onto a subgroup of TuBu neurons (TuBus) located in the AOTU neuropil. These anterior-projecting DNs (APDNs) also activate TuBu neuronal activity, which project specifically to the superior bulb region (BU_sup_). These neurons are functionally connected in turn to EB ring neurons (EB-R2/R4m) of the central complex, which are involved in sleep regulation. Interestingly, activation of the APDNs can induce calcium oscillations in the cell bodies of EB ring neurons as well as their axonal ring structure. Blocking neurotransmission within this circuit affects sleep and causes a dramatic reduction in the sleep-wake arousal threshold. The data together show that this APDN-TuBu_sup_-EB circuit regulates sleep drive and the sleep-wake arousal threshold in adult *Drosophila*.

## Results

### Intersectional strategy assesses expression and function of circadian neurons

To investigate the circadian circuit neurons important for sleep- and wake-regulation, we performed an activation screen. Because there are only a handful of GAL4 drivers that exclusively label circadian neurons subsets, we used an intersectional strategy that included the classical circadian driver *Clk856-GAL4* (Figure 1a). Our immunohistochemistry results, as well as previous studies (Gummadova et al., 2009), confirmed that this driver expresses in most central circadian neurons in adult flies (Figure 1b left panel). To characterize the expression patterns and behavioral output of other circadian neuron drivers, we made a stable line (*Clk856-GAL4, LexAop-FLP; UAS-FRT-stop-FRT-CsChrimson.mVenus*) to cross with different LexA lines from the FlyLight library collection (Pfeiffer et al., 2008). The LexA drives the expression of flippase to excise the stop codon, which will then activate the expression of functional red-shifted and YFP-tagged channelrhodopsin-CsChrimson-mVenus (Klapoetke et al., 2014), only within the intersectional territory (Figure 1a). Experimental and control flies were loaded into 96-well plates containing ATR food. After 3 days of baseline sleep recording, different subsets of circadian neurons were activated for 24-hours with pulsed red light. Sleep and activity were monitored by Flybox automated video-recording (Guo et al., 2017).

**Figure 1.**
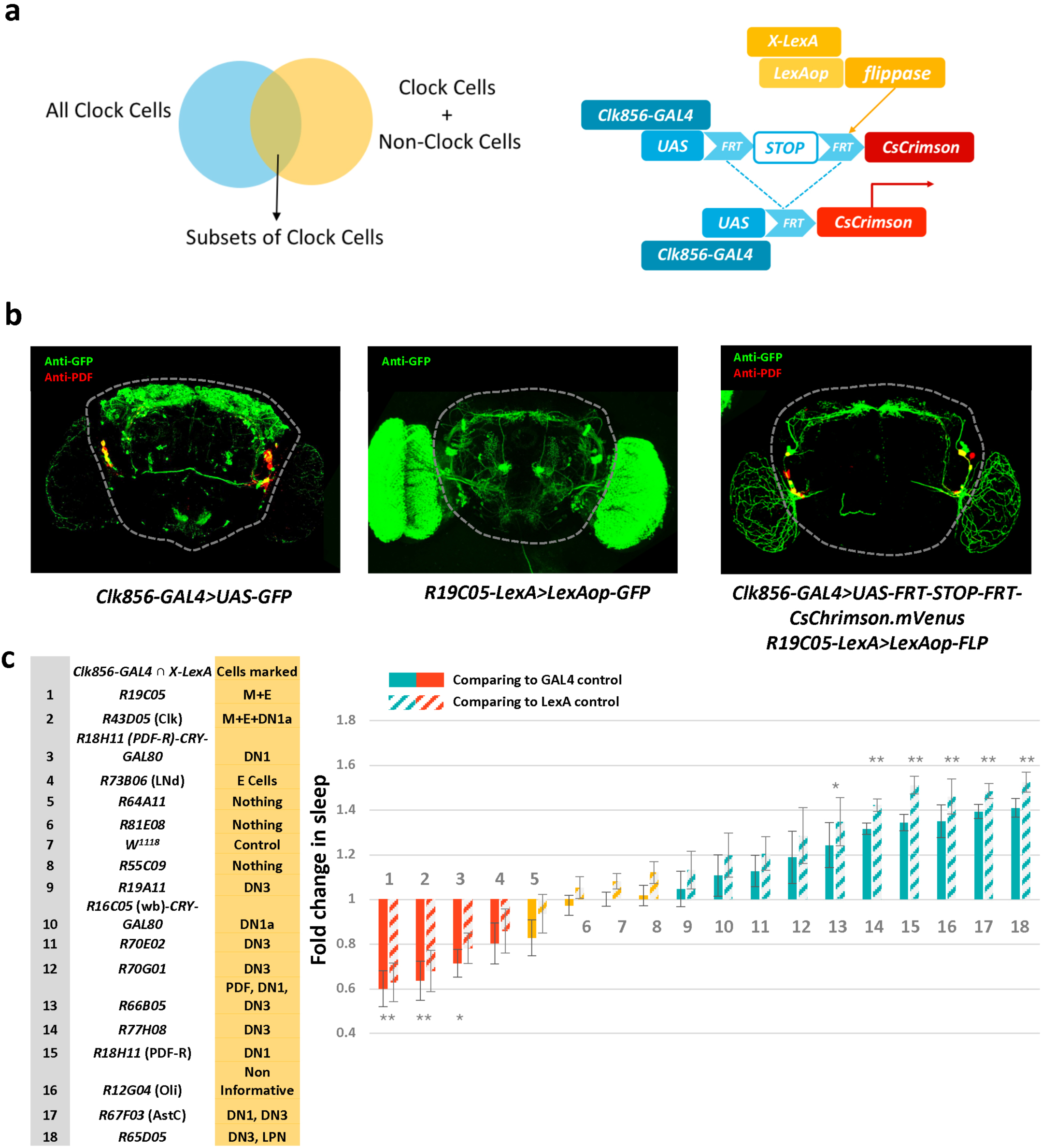
Intersectional strategy to assess behavioral phenotypes caused by activation of different circadian neuron subsets. **a,** Strategy to label only subsets of clock cells to test gain of function behavior of sleep and activity assay. Functional CsChrimson.mVenus is expressed in clock neurons by *Clk856-GAL4* where in the flippase is expressed by *LexA* to excise the stop codon. **b,** Example to show the intersection strategy. Left panel, anti-GFP labelling all the clock cells expressing the *Clk856-GAL4*. Middle panel, expression pattern of *R19C05-LexA* from Flylight. Right panel, anti-GFP labelling the E-cells (LNds) and the M-cells (lLNvs and sLNvs), which are the common cells between the *Clk856-GAL4* and the *R19C05-LexA* drivers. **c,** Total fold changes of sleep in experimental groups comparing to parental controls. Red histograms indicate activity-promoting groups and the blue bars indicate the sleep-promoting groups compared to parental controls. Solid bars indicate sleep changes comparing to *Clk856-Gal4>UAS-FRT-STOP-FRT-CsChrimson, LexAop-flippase/+*) and striped bars indicate sleep changes comparing to *X-LexA/+*. n=8-12 in each group. All experiments were repeated three times except for *Clk856-GAL4 ∩ R18H11-LexA* and *Clk856-GAL4 ∩ R67F03-LexA* groups (four repeats). Data are presented as mean ± SEM; *p < 0.05, **p < 0.001 are significant differences from the control group (one way ANOVA with Tukey post hoc test)

Acute activation of different circadian neurons triggers different behavioral responses (Figure 1c). A persistent wake-promoting/sleep-reducing effect during the LED stimulation period occurs whenever the expression pattern includes E-cells, consistent with our previous study that E cells are the major activity-promoting circadian neurons (Guo et al., 2014; Guo et al., 2017). In contrast, activating subsets of the DN1s and DN3s significantly increases sleep (Figure 1c). The *Clk856-GAL4 ∩ R65D05-LexA* line expresses in Ast-A positive circadian neurons and had the strongest sleep-promoting effect. This phenotype is consistent with a recent report characterizing the important role of Ast-A-expressing neurons in sleep regulation (Chen et al., 2016). The remaining available LexA lines also labeled sleep-promoting circadian neurons and show selective and consistent expression within the DN1s and DN3s (see below), indicating that these two classes of circadian neurons contribute to one or more sleep-promoting circadian circuits.

### Distinct circadian neuron output circuits for sleep and activity regulation

To better understand the detailed organization of these activity and sleep-promoting circadian neurons, we performed double immunostaining with anti-PDF and anti-GFP on two strong sleep-promoting lines: *Clk856-GAL4 ∩ R18H11-LexA* and *Clk856-GAL4 ∩ R67F03-LexA* (Figure 2a and 2b). Both lines clearly and consistently label subsets of circadian DN1s and DN3s (Figure S1). *R18H11* is a traditional circadian driver line derived from the *Pdfr* gene, and *R67F03* is derived from the *Ast-C* gene (Jenett et al., 2012). These neurons have posterior branches that overlap with dorsal projections of sLNvs (M cells) and LNds (E cells), consistent with previous results indicating that sleep-promoting DN1s inhibit wake-promoting M and E cells. In addition, both DN1s and DN3s have major branches that spread into the anterior brain region called the SLP (superior lateral protocerebrum) (Supplemental Video 1), hence the name anterior-projecting DNs (APDNs).

**Figure 2.**
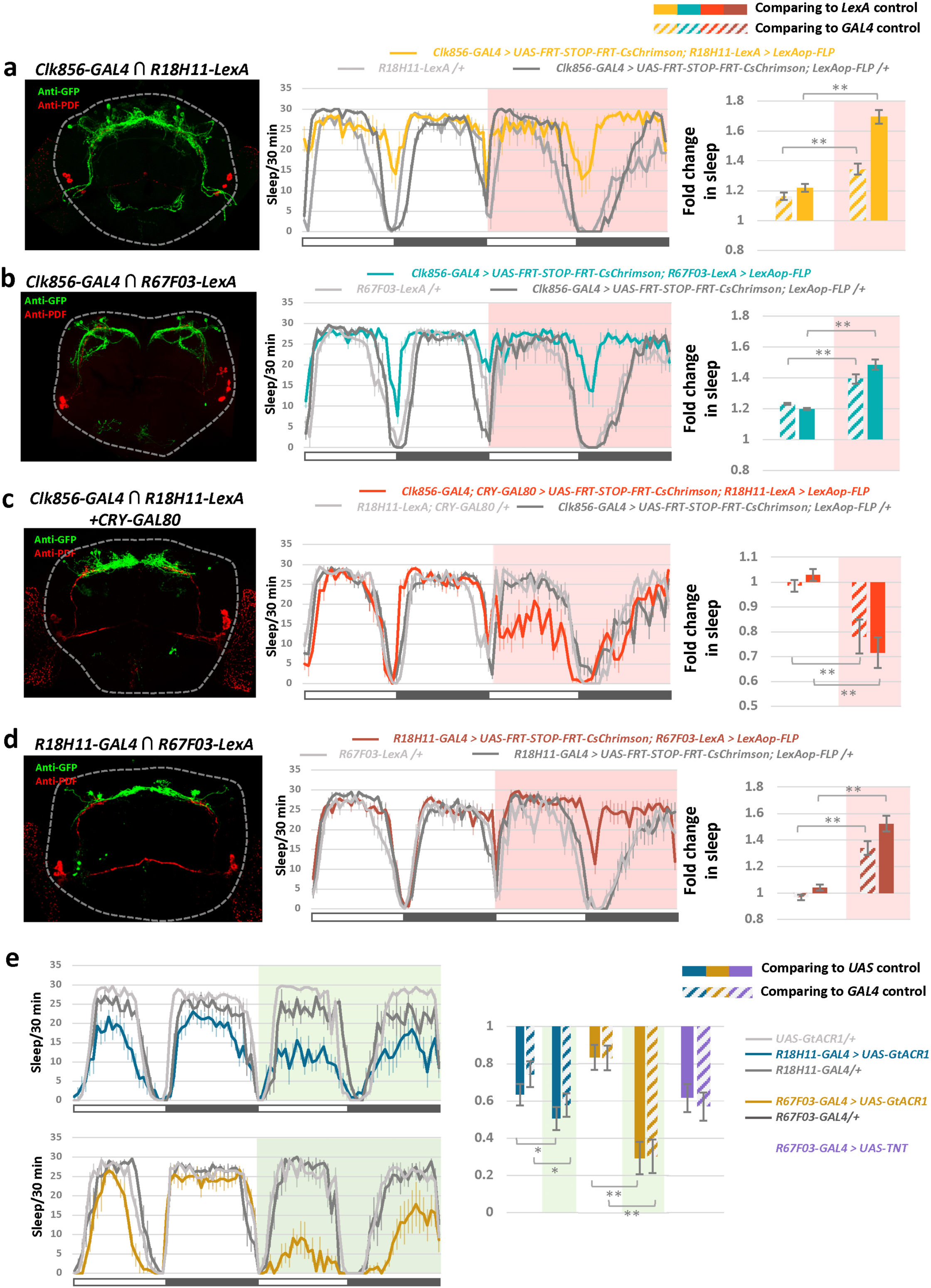
DN1s consist of two anatomically and functionally sets of neurons. **a-d,** Left panel, anti-GFP and anti-PDF visualizing the CsChrimson.mVenus expression within clock cells common between the *Clk856-GAL4* and the chosen *LexA* lines. (**a**) The cells are the sleep-promoting APDNs including DN1s and DN3s as well as PI-projecting DN1s; (**b**) sleep-promoting APDNs including DN1s and DN3s; (**c**) CRY negative, PI-projecting, activity-promoting DN1s; (**d**) sleep-promoting DN1s. Middle panel, sleep curves of experimental and control flies in baseline day and red LED day. The genotypes for control and experimental groups are labeled above the panel. Right panel, quantification of relative fold change of sleep on both control day and LED day (pink background). Stripe color bars indicate sleep from experimental group divided by GAL4 control and solid color bars indicate sleep from experimental group divided by LexA control, n=12-16 for experimental groups and n=8 for control groups. Data are presented as mean ± SEM; **p < 0.001 are significant differences from both control group in LED day and the experimental groups in baselineday by one way ANOVA with Tukey post hoc test. **e,** Left panel, sleep curves of *R18H11-GAL4>UAS-GtACR1* and *R67F03-GAL4>UAS-GtACR1* as well as control flies in baseline day and green LED day. Right panel, quantification of relative sleep fold change on the baseline day and green LED day (green background) of *R18H11-GAL4>UAS-GtACR1, R67F03-GAL4>UAS-GtACR1* and *R67F03-GAL4>UAS-TNT* with their respective controls, Stripe color bars indicate sleep from experimental group divided by GAL4 control and solid color bars indicate sleep from experimental group divided by UAS control n=16 for experimental groups and n=8 for control groups, ‘*’ indicates p < 0.05 and ‘**’ indicates p < 0.001 by one-way ANOVA with Tukey’s post-hoc test. Error bars correspond to SEM. All experiments were repeated three times.

Activation of those APDNs strongly promotes sleep in the LED day compared to controls (Figure 2a-b). However, some sleep-promoting effects of the APDNs was also observed during the baseline day. This may be due to entrainment light activation of CsChrimson expressed in DN3s. These neurons are located closer to the surface of brain, and there was no baseline day effect when CsChrimson was expressed only in DN1s (Figure 2d).

Despite the similarities of these anterior-projecting branches, the two sets of DNs have distinct morphological differences around the PI region, a functional homolog of the mammalian hypothalamus (de Velasco et al., 2007). *Clk856-GAL4 ∩ R18H11-LexA* DNs form a tight neuropil in the PI region (Figure 2a), whereas *Clk856-GAL4 ∩ R67F03-LexA* DNs barely have fibers near the PI (Figure 2b and Figure S1). To address this distinction, we added a *CRY-GAL80* transgene to the *Clk856-GAL4 ∩ R18H11-LexA* combination, which identified a CRY-negative (cryptochrome-negative) subset of DN1s that appear to send projections only to the PI (Figure 2c). Activation of these PI-projecting DN1s promoted wakefulness and inhibited sleep (Figure 2c), consistent with previous reports that PI-projecting DN1s connect to DH44+ PI neurons to regulate locomotor activity (Cavanaugh et al., 2014; Kunst et al., 2014).

We next examined projections of the R18H11 sleep-promoting DN1s. To access these processes and to also examine whether the R67F03 and R18H11 drivers label some sleep-promoting DN1s in common, we identified their overlapping neurons by combining *R67F03-LexA* and *R18H11-GAL4*. This strategy identified 4-5 DN1s which have clear posterior and anterior fibers (Figure 2d) and strongly promote sleep. These results indicate that DN1s are heterogeneous with distinct morphological patterns and behavioral contributions.

To further confirm the contribution of these APDNs to regulate sleep under basal conditions, we temporally hyperpolarized these neurons by expressing the cyan-gated anion channel GtACR1 and exposed adult flies to green light after 3 days of ATR feeding (Mohammad et al., 2017). This caused a significant sleep reduction in both *R18H11>GtACR1* and *R67F03>GtACR1* flies during the green LED day (Figure 2e), consistent with our previous result that these APDNs promote sleep. We note that *R18H11>GtACR1* flies also sleep less compared to controls during the baseline day, suggesting that some GtACR1-expressing *R18H11-GAL4* DN1s may be inhibited by the FlyBox entrainment light.

To address whether these neurons promote sleep through neurotransmitter or neuropeptide secretion, we expressed TNT (tetanus toxin light chain) because it specifically inactivates synaptobrevin (Sweeney et al., 1995), which is necessary for classical neurotransmitter release. *R67F03>TNT* flies significantly reduced total sleep amounts and dramatically increased locomotor activity during day and night (Figure 2e and data not shown). The data taken together demonstrate an important role for the APDN circuit in sleep regulation.

### Anterior-projecting DN1s innervate the AOTU neuropil

To examine whether the anterior fibers from the APDNs are receiving or sending signals, we labelled brains with the post-synaptic (dendritic) and pre-synaptic (axonal) fluorescent markers Denmark (Figure 3a-red) and syt1-GFP (Figure 3a-green) respectively (Nicolai et al., 2010). First, the posterior projections of DN1s are presynaptic as well as postsynaptic; they descend into the accessory medulla, where they reach the sLNv cell bodies (Figure 3a middle panel). DN1 fibers have been previously shown to connect to the dorsal branches of lateral M and E cells, and these data suggest that the DN1s receive as well as send signals from/to the lateral M and E pacemakers (Guo et al., 2016). Second and not unexpectedly, presynaptic markers densely label the DN1 projection to the PI region, consistent with the evidence that some activity promoting DN1s output to PI neurons (see above). Third and most importantly, the DN1 anterior branches are solely and robustly labelled by the pre-synaptic marker, indicating that the neuropil in SLP region is a new major post-synaptic APDN target (Figure S2).

**Figure 3.**
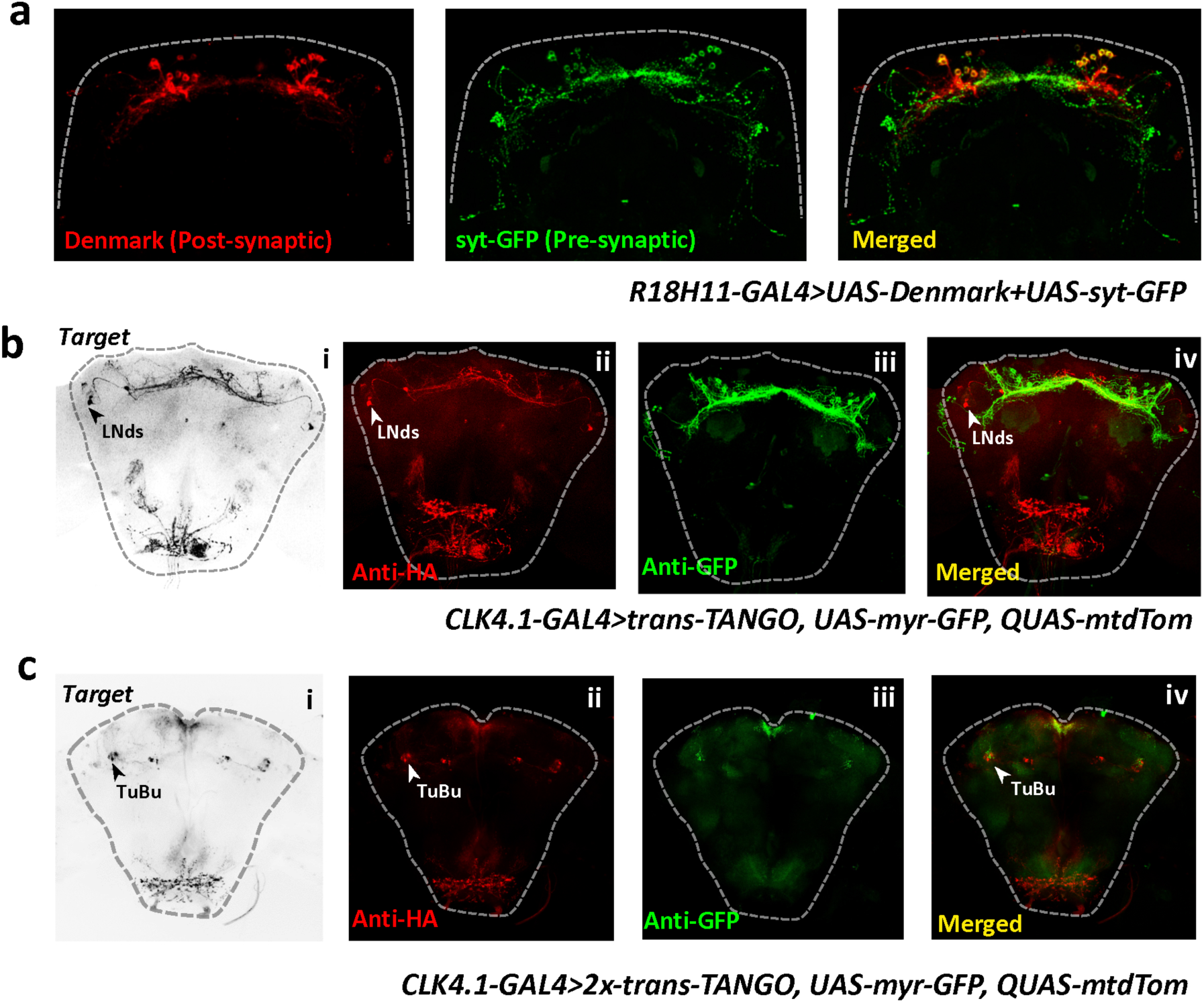
APDNs form diverse output synapses in adult *Drosophila*. **a,** *R18H11-GAL4>UAS-Denmark+UAS-synaptotagmin-GFP* brains were dissected and stained with anti-DsRed to identify the DN1s dendritic regions (red; left panel). To identify the synaptic regions of DN1s, the brains were stained with anti-GFP (green; middle panel). These patterns were aligned and overlaid (merged panel on the right). **b,** Trans-synaptic labelling using *trans*-Tango. Brains of *CLK4.1M-GAL4>trans-Tango, UAS-myr-GFP, QUAS-mtdTom* flies were dissected and stained with anti-HA to identify the DN1 post-synaptic partners (**i-**black, **ii-**red) - E-cells (LNds) are downstream of DN1s. The DN1s were visualized with GFP antibody staining (**iii**-green). These patterns were aligned and overlaid (**iv**). **c**, To better visualize the downstream partners, two copies of the *trans*-Tango were expressed in the DN1s. Only the anterior sections of the brain are shown to visualize TuBu neurons. Brains of *CLK4.1M-GAL4>2x-trans-Tango, UAS-myr-GFP, QUAS-mtdTom* flies were dissected and stained with anti-HA to identify the DN1 post-synaptic partners (**i**-black, **ii**-red) – a subset of TuBu neurons are marked as downstream of DN1s. The DN1 anterior fibers were stained with anti-GFP (**iii**-green). These patterns were aligned and overlaid (**iv**).

To trace and define the specific downstream synaptic partners located in the SLP, we employed the new developed anterograde trans-synaptic tracing tool *trans*-Tango (Talay et al., 2017). Expression of the tethered *trans-*Tango ligand in neurons can trigger mtdTomato expression in post-synaptic targets. Therefore, we co-expressed the *trans-*Tango ligand and myrGFP in DN1s to visualize presynaptic neurons. There was clear postsynaptic mtdTomato labeling in several clusters of neurons within the circadian circuit, including the LNds and some DN3s (Figure 3b and data not shown). The LNd (E cell) output fits with previous data (Guo et al., 2016). The DN3 output indicates that these neurons are new post-synaptic targets of DN1s (data not shown; see Discussion).

Because we could not detect any postsynaptic signal close to the two additional major contact sites of DN1s, the PI and SLP regions, we expressed two copies of the *trans*-Tango ligand in DN1s, which then resulted in positive trans-synaptic signal in both locations (Figure 3c). In the SLP, the myrGFP-labeled anterior projections of the APDNs reach the surface and innervate a neuropil called the anterior optic tubercle (AOTU). The postsynaptic mtdTomato signal is expressed in a subset of tubercular-bulbar (TuBu) neurons not previously associated with the circadian circuit (Figure 3c). These neurons are reported to receive visual information from the AOTU and send it into the bulb (BU) neuropil (Omoto et al., 2017), where they connect to the ellipsoid body (EB). EB neurons are essential for a variety of functions, including sleep homeostasis and arousal (Lebestky et al., 2009; Liu et al., 2016). Thus, we hypothesized that the sleep-promoting APDNs send circadian information to EB neurons to regulate sleep.

To enhance the details of the connections between the APDNs and TuBus within the dense AOTU neuropil, we labeled the two groups of neurons with different membrane-tethered fluorescence proteins using (*R18H11-LexA>LexAop2-mCD8::GFP; R76B06-GAL4>UAS-IVS-mCD8::RFP*). Co-staining verified that the anterior neurites from APDNs innervate dendrites of a subset TuBus (Figure 4a).

**Figure 4.**
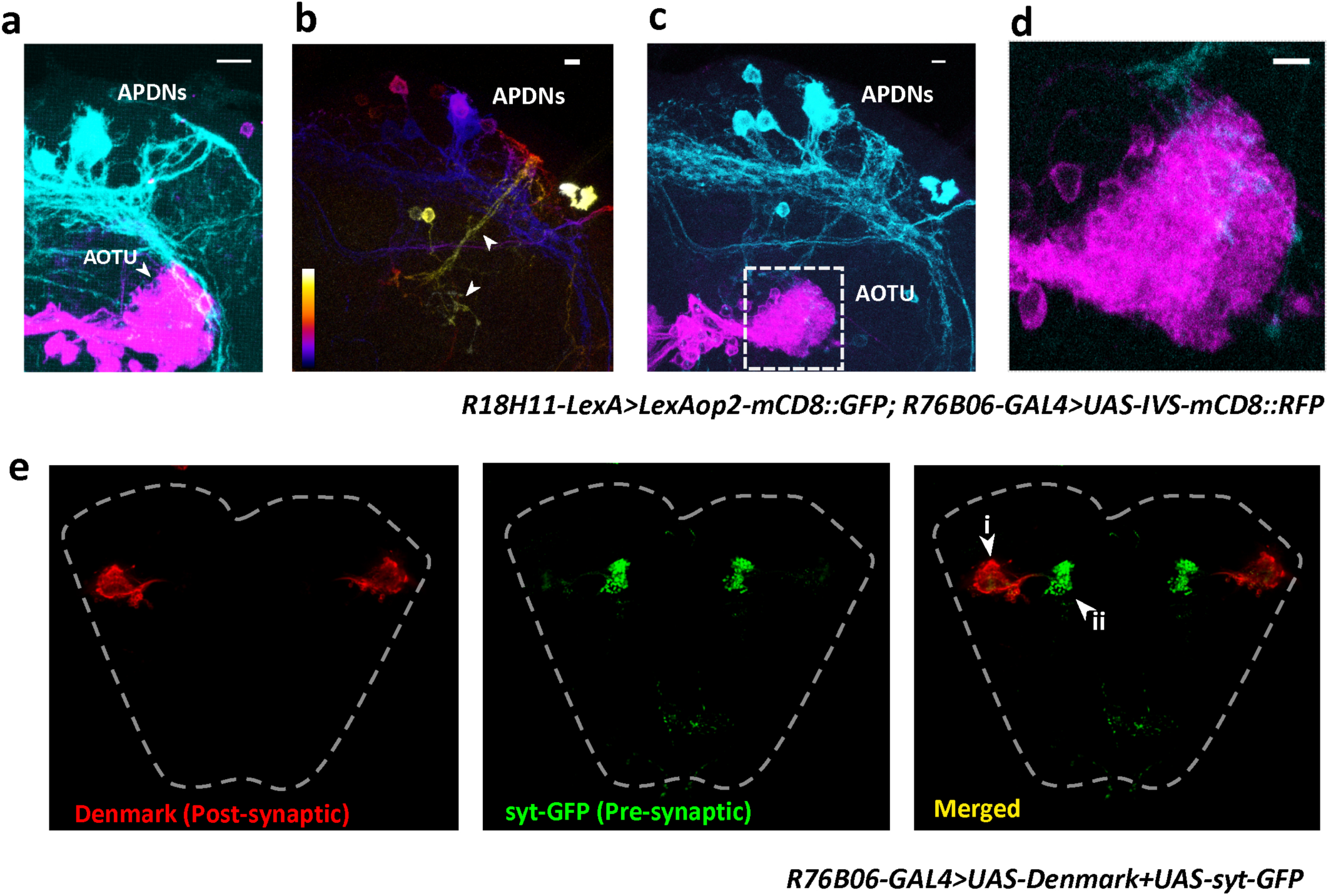
Sleep-promoting APDNs connect to the AOTU neuropil. **a,** Maximum projection of whole mount brain of *R18H11-LexA>LexAop2-mCD8::GFP; R76B06-GAL4>UAS-IVS-mCD8::RFP* fly after regular IHC. (R18H11-cyan, R76B06-magenta) **b,** Maximum projection of whole mount brain after ExM showing z-depth of DN1s. Blue indicates most posterior, white/yellow indicates most anterior of the stack (slices #1-206). **c,** Maximum projection of whole mount brain after ExM of the complete circuit. **d,** Zoomed in view of boxed region of Scale bar= 10um for all. **e,** *R76B06-GAL4>UAS-Denmark/ UAS-synaptotagmin-GFP* brains were dissected and stained with anti-DsRed to identify the TuBu dendritic regions (red; left panel). The synaptic regions of TuBu were stained with anti-GFP (green; middle panel). These patterns were aligned and overlaid (merged panel on the right, arrow **i** indicates dendrites and arrow **ii** indicates synapses).

Protein-retention expansion microscopy (pro-ExM) was also exploited to investigate in more detail the overlapping pattern inside the AOTU (Chen et al., 2015). Post-ExM confocal images with z-depth color coding show the elaborate structure of the DNs, specifically how far the APDN anterior fibers project (Figure 4b, yellow spectrum in middle panel) and their extent of overlap with the AOTU neuropil (Figure 4c and d). Moreover, these APDN fibers are anatomically restricted to specific sub-classes of the AOTU. To verify that TuBus receive input from anterior projections of APDNs, we used the broader TuBu driver *R76B06-GAL4* to express syt1-GFP and Denmark; *R76B06-GAL4* expresses in different classes of TuBus. Indeed, TuBus have dendrites in the AOTU neuropil (Figure 4e i) and presynaptic boutons in the BU region; this is where EB ring neurons send dendrites (Figure 4e ii), suggesting that APDNs signal to EB neurons through TuBus.

### APDNs activate the BU-projecting TuBu neurons to promote sleep

TuBu neurons are involved in processing visual information into the EB ring neurons, and different classes of TuBus are organized to output in parallel to different parts of the BU (Omoto et al., 2017). To address functional connections between the anterior projections of APDNs and these different TuBus, we expressed the ATP-activated cation channel P2X2 in DN1s with *R18H11-LexA* and expressed the genetically encoded calcium indicator GCaMP6f under the control of TuBu driver *R76B06-GAL4*. After baseline recording, ATP was added to evoke firing in the APDNs while monitoring the TuBus. A sparse subset of TuBus exhibited consistent and significant calcium increases in response to the DN1 activation (Figure 5a and Supplemental Video 2).

**Figure 5.**
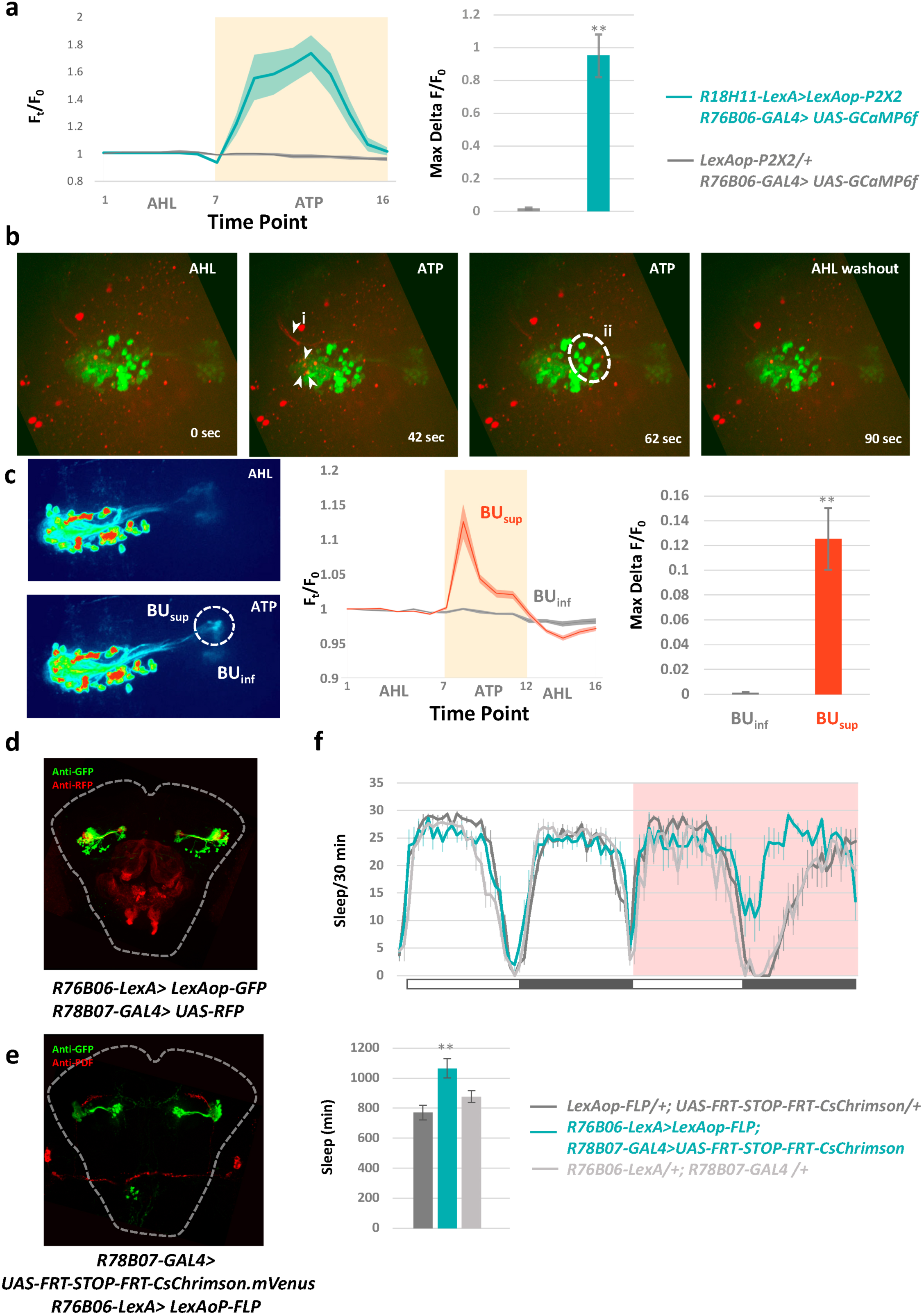
APDNs activate superior-projecting TuBu (TuBu_sup_) neurons to promote sleep. **a,** Left panel, relative fold changes of calcium level in *R18H11-LexA>LexAop-P2X2; R76B06-GAL4>UAS-GCaMP6f* fly brain during perfusion. The quantification of peak GCaMP6f changes is plotted in the right panel. n=5. ‘**’ indicates p<0.001 by student’s t-test. Shading and error bars correspond to SEM. **b,** Sequence of frames when *R18H11-LexA> LexAop-P2X2, LexAop-RCAMP; R76B06-GAL4>UAS-GCaMP6f* brain is perfused with ATP. i- Increase in RCaMP signal observed during ATP perfusion neurons in DN1 anterior fiber as well as in the AOTU neuropil where DN1s send terminals (arrows). ii- Increase in GCaMP levels in the TuBu neurons cell bodies during ATP perfusion due to DN1 activation (white dashed circle). **c,** Activation of APDNs induce calcium increase in the superior branch of the TuBu neurons specifically (left panel). Middle and right panels, quantification of the increase in GCaMP signal in the superior and the inferior branches of the TuBu neurons. The quantification of maximum GCaMP6f changes is plotted in the right panel. n=3. ‘**’ indicates p<0.001 by student’s t-test. Shading and error bars correspond to SEM. **d,** *R78B07-GAL4* (Anti-RFP) labelled subset of TuBu neurons which are overlap with the *R76B06-LexA*(Anti-GFP) labelled TuBu_sup_. **e,** Anti-GFP labelling the CsChrimson.mVenus expressing cells common between the *R78B07-GAL4* and the *R76B06-LexA*. The cells marked here are the TuBu_sup_. **f,** Top panel, sleep curves of experimental and control flies in baseline day and LED day. Bottom panel, quantification of total sleep on the LED-day. The genotypes for each group were labelled next to the panel. n=16 for each group, Error bars correspond to SEM. ‘**’ indicates p<0.001 by one-way ANOVA with Tukey’s post-hoc test.

To verify this excitatory input, we performed a two-color calcium imaging experiment with two independent binary expression systems: the red-shifted calcium indicators R-CaMP6 and P2X2 were expressed in APDNs, and GCaMP6f was expressed in the TuBus. The two calcium indicators have different excitation/emission spectra. The ATP-evoked calcium influx can be seen to spread from DN1 cell bodies to the anterior fibers, followed by a calcium increase in several presynaptic terminal boutons of the AOTU neuropil. A subset of TuBus was then triggered (Figure 5b and Supplemental Video 3). The results indicate that sleep-promoting APDNs induce firing of a subset of downstream TuBus.

TuBus are comprised of different subclasses with axon projections to different Bus: superior (BU_sup_), inferior BU (BU_inf_) and anterior BU (BU_ant_) (Lovick et al., 2017). Moreover, the different TuBus connect to different EB ring neurons (Omoto et al., 2017). To trace where the TuBus transmit information after APDN firing, we recorded the calcium response in different BU regions. Interestingly, we only observed calcium influx spreading to BU_sup_ during ATP perfusion period in *APDN>P2X2, TuBu>GCaMP6f* flies (Figure 5c). We then asked whether this subset of TuBus can promote sleep by identifying *R78B07-GAL4* as specifically targeting the specific BU_sup_ -projecting TuBus (TuBu_sup_) (Figure 5d). Because this GAL4 also drives ectopic expression beyond the TuBu_sup_ (Figure 5d), we used the intersectional strategy to restricted the CsChrimson expression to TuBu_sup_ (*R78B07-GAL4, R76B06-LexA >LexAop-FLP, UAS-FRT-stop-FRT-CsChrimson*). Immunohistochemistry indeed confirms that CsChrimson expression is only in TuBu_sup_ (Figure 5e).

During the baseline day (without LED stimulation), sleep levels are indistinguishable between experimental flies and parental controls, but stimulation of TuBu_sup_ with 24-hr red LED illumination strongly inhibited locomotor activity and increased sleep compared to the genetic controls. (Figure 5f). The results taken together identify a novel sleep-promoting circuit, from circadian APDNs to specific TuBu_sup_.

### The APDN-TuBu-EB circuit regulates sleep-wake arousal threshold

TuBu neurons connect to distinct EB ring neurons via different BU_sup_, and individual layers of EB ring neurons can play unique roles in regulating different behaviors. For instance, R2/R4m neurons encode sleep pressure as well as gate mechanical stress-induced arousal responses (Lebestky et al., 2009; Liu et al., 2016). We therefore asked, are EB-R2/R4m neurons post-synaptic targets of TuBus? If so, are they the only EB ring neuron targets and what kind of input signal is conveyed to the EB from TuBus? To answer these questions, we expressed GCaMP3 instead of mtdTomato under control of the *trans-*Tango tracing tool, while co-expressing the *trans*-Tango ligand and P2X2 cation channel with the TuBu driver *R76B06-GAL4*. This transynaptic “label and activate” strategy not only screens pan-neuronally to identify post-synaptic neurons, but also specifies the direction (activation or inhibition) between pre- and post-synaptic partners (Figure 6a).

**Figure 6.**
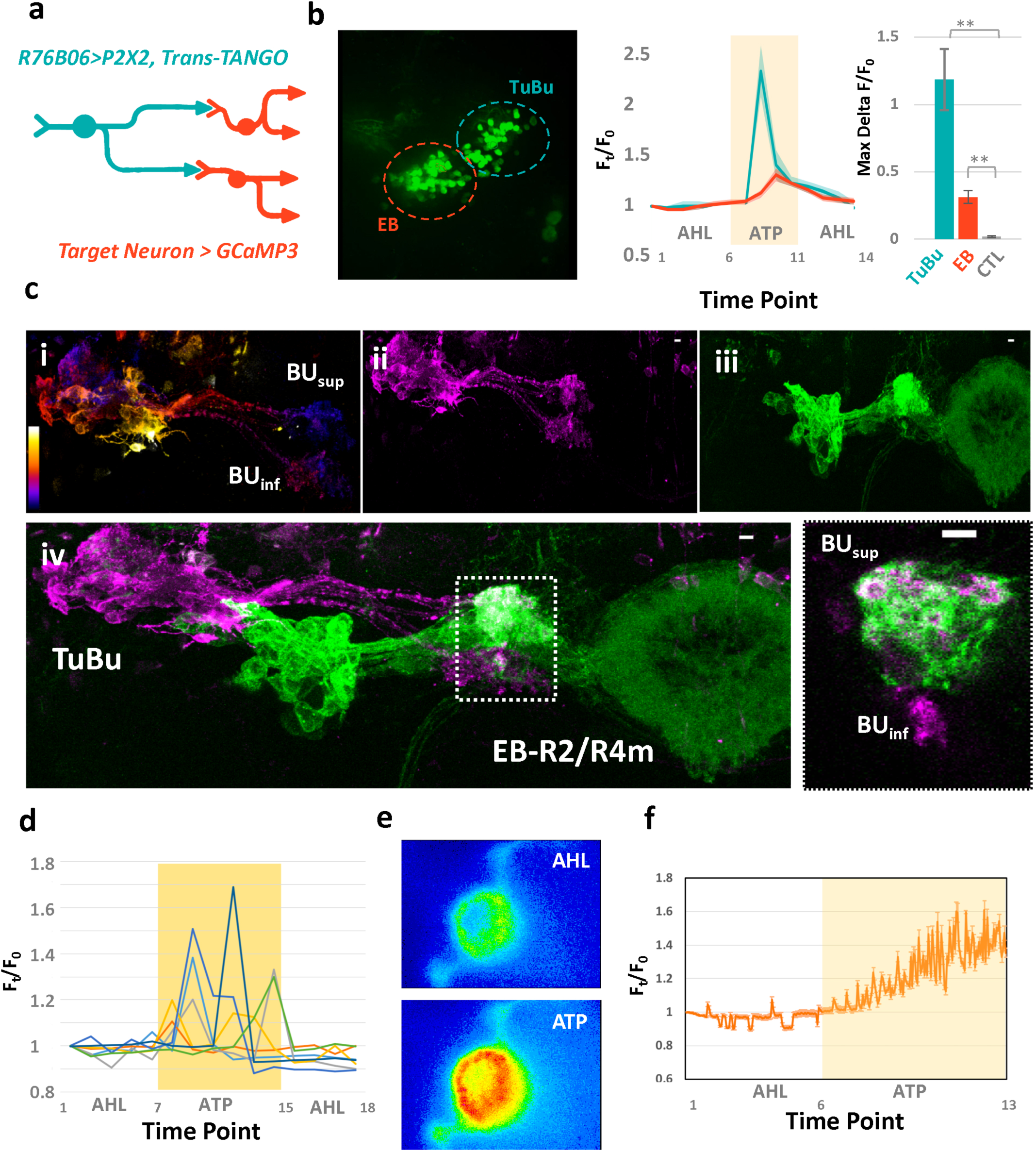
APDNs promote calcium oscillations in the EB ring neurons via TuBu_sup_ neurons. **a,** Strategy to label the downstream neurons and assay the connection by combining *trans-*Tango with P2X2 and GCaMP3. **b,** Left panel, GCaMP3 expression driven by *R76B06-GAL4>trans-*Tango labels both TuBu neurons and the downstream EB ring neurons. Middle panel, relative fold changes of calcium level in *trans-*Tango*/UAS-P2X2; R76B06-GAL4/QUAS-GCaMP3* fly brain during perfusion. Right panel, quantification of peak calcium changes in *trans-*Tango*/UAS-P2X2; R76B06-GAL4/QUAS-GCaMP3* flies. n=3, Shading and error bars correspond to SEM. **c,** ExM images from representative *R32H08-LexA>LexAop2-mCD8::GFP; R76B06-GAL4>UAS-IVS-mCD8::RFP* fly brain (i) Maximum projection of whole mount brain after ExM showing z-depth of TuBu neurons as well as BU_sup_- and BU_sup_-projections. Blue indicates most posterior, white/yellow indicates most anterior of the stack. (ii) Maximum projection of TuBu channel only (magenta). (iii) Maximum projection of EB channel only (green). (iv) Maximum projection showing merge of TuBu and EB. Right panel, single focal plane of boxed region to show significant overlap. **d,** Activation of APDNs using *R18H11-LexA> LexAop-P2X2* increases calcium oscillations in the EB ring cell bodies measured using GCaMP6f (*R41A08-GAL4>UAS-GCaMP6f*). Different colored traces indicate calcium response from individual EB ring neurons (cell bodies). **e,** Heat-map showing increase in calcium oscillations in EB ring structure before (upper panel) and after (lower panel) APDNs are activated in *R18H11-LexA>LexAop-P2X2, VT038828-GAL4>UAS-GCaMP6f* flies. **f,** A representative real-time GCaMP6f response trace in *R18H11-LexA> LexAop-P2X2, VT038828-GAL4>UAS-GCaMP6f* flies.

Indeed, we found that some EB ring neurons are labeled by *R76B06> trans-Tango* flies. Since TuBus labeled by this driver project to both BU_sup_ and BU_inf_, they must connect to different ring neurons (Figure S3). The *R76B06>trans-Tango* also labels the TuBus themselves. This could be due to the interaction of *trans-*Tango ligand and receptor expressed within neighboring TuBus (Figure 6b). Moreover, ATP-evoked TuBu activation led to a subsequent modest but significant calcium increase in EB ring neurons (Figure 6b and Supplemental Video 4), consistent with a previous report (Sun et al., 2017) and indicating that the EB ring neurons receive information from the APDNs via TuBus.

To address which ring neurons receive stimuli from TuBu_sup_, we used high-resolution ExM imaging and confirmed that EB-R2/R4m neurons labeled by the *R32H08-LexA* show potent overlap with the TuBu_sup_ (Figure 6c) (Xie et al., 2017), other layers of EB ring neurons barely connect to that part of BU_sup_ (data not shown). The data suggest that APDNs specifically activate TuBu_sup_ and then input to EB-R2/R4m neurons, which are essential for sleep and arousal regulation.

To verify this input pathway, we activated APDNs with ATP perfusion in *R18H11>P2X2* flies and assayed the change in EB ring neurons by expressing the calcium indicator GCaMP6f with different EB ring neuron drivers. As expected, we detected an increased calcium response in the cell bodies of these defined EB ring neurons during the period of ATP perfusion. Surprisingly, ATP-evoked APDN excitation also triggers a strong increase in calcium oscillations in these EB ring neurons compared to the more modest calcium increase in EB ring neurons caused by activation of TuBus (Figure 6d and Supplemental Video 5). These fluctuations are coordinated in an alternating fashion within distinct subclasses of EB ring neurons, which presumably reflects the connection patterns between different EB neurons (Figure 6d). The APDN activation also promotes long-lasting and intense high-frequency calcium oscillations in the axonal ring structure, indicating that EB ring region neuronal activity is sensitive to APDN input (Figure 6e-f and Supplemental Video 6). This result suggests that EB neuronal activity is under circadian control and that the EB-R2/R4m sleep homeostatic center receives circadian inputs to regulate sleep levels.

To examine the epistatic interaction of APDNs and EB-R2/R4m neurons in basal sleep and circadian regulation, we used two binary systems to simultaneously express CsChrimson in APDNs and GtACR1 in EB ring neurons (see Figure 7 legend). Experimental and control flies were fed with ATR and entrained with 12:12 LD dim white light in a FlyBox for 3 days. The flies were then exposed to a mechanical stimulus every 30 min as well as illuminated with both 630nm and 540nm LEDs on day 4. The strategy was to activate APDNs while hyperpolarizing EB ring neurons (Figure 7a).

**Figure 7.**
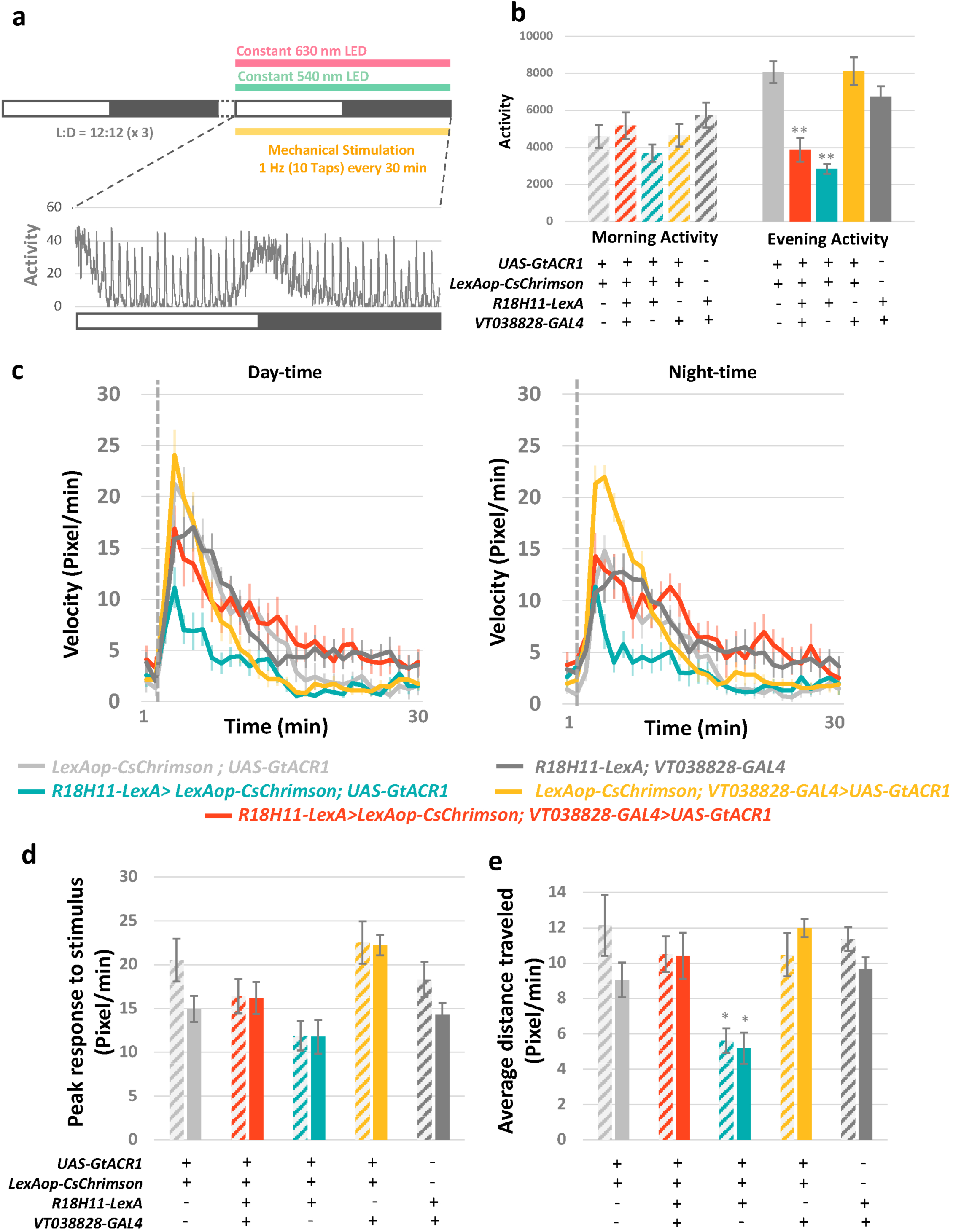
The APDN-EB circuit regulates sleep-wake arousal threshold. **a,** Experimental setup to test sleep-wake test arousal threshold in flies using the Flybox. The flies were entrained for 3 days in a 12:12hr LD cycle. On the 4th day, the flies were exposed to constant green LED (540 nm) and red LED (630nm) and were also mechanically stimulated for 24 hours with 10 taps (1HZ) every 30 minutes. A representative activity plot from a control group was plotted bellow. **b,** Morning (ZT0-6) and evening activity (ZT11-15) in different groups. The genotypes for each group were labelled bellow. n=8 for each group. Error bars correspond to SEM. ‘**’ indicates p<0.001 by one-way ANOVA with Tukey’s post-hoc test. **c,** Average response to mechanical stimulation during day-time (ZT7-11) and night-time sleep period (ZT19-23) in different groups. Genotypes for each group were labelled bellow. Data from 10 repeats were calculated to plot the velocity traces for each groups. Gray dashed lines indicate mechanical stimulation. Error bars correspond to SEM. **d, e,** Peak response to stimulus (**d**) and average distance travelled in 15min after stimulus (**e**). Genotypes are shown below the data. n=10 for each groups. Error bars correspond to SEM. ‘**’ indicates p<0.001 by one-way ANOVA with Tukey’s post-hoc test.

It is notable that activation of APDNs inhibits locomotor activity and promotes sleep during the evening (Figure 7b-teal bar), consistent with the notion that DN1s inhibit activity-promoting E-cells. Moreover, this phenotype was not affected when EB ring neurons were simultaneously silenced (Figure 7b-red bar), suggesting that the anterior output circuit does not interfere with the locomotor-inhibiting role of the APDN posterior projections.

We next focused on the mechanical induced arousal response during the siesta and nighttime sleep periods when flies exhibit little baseline activity. All groups show a period of locomotor hyperactivity after a strong stimulation, and APDN-activated flies display a slightly reduced but comparable startle peak response, indicating that their mobility is not markedly affected by the ring neuron inhibition (Figure 7c and d). However, these flies fail to maintain prolonged wakefulness after the startle response as evidenced by the distance traveled during the post-stimulation period; it is significantly lower than the other groups (Figure 7e-teal bar). The quick decline in locomotor activity of *APDN>CsChrimson* flies is due to activation of the APDN-TuBu_sup_-EB circuit since silencing EB ring neurons can rescue the startle-induced locomotor hyperactivities (Figure 7c and e-red bar). These data demonstrate that this circuit plays an important role in regulating arousal levels and suggest the APDN-induced calcium oscillations in this sleep homeostasis center reduces the response to sensory input to facilitate the wake/sleep switch.

## Discussion

Sleep in diverse systems is characterized by a set of common features. They include reduced locomotor activity, increased arousal threshold and altered neuronal oscillation patterns in higher brain centers (Yap et al., 2017). Although there is no single mechanism that can account for all these features, we present here a novel anatomical circuit in the fly brain that influences all of them. The circuit couples two important centers: the EB ring neurons, which determine sleep drive and arousal levels (Lebestky et al., 2009; Liu et al., 2016); and the circadian APDN1s, which contribute to the daily sleep/wake pattern (Guo et al., 2016).

The DN1s have been previously reported to be both sleep promoting (activity-inhibiting) and activity promoting (sleep-inhibiting) (Cavanaugh et al., 2014; Guo et al., 2016; Kunst et al., 2014), a complication that has not plagued other circadian neuron groups. For this reason, we began this study with an intersectional activation and anatomical screen of specific circadian neuron subsets. The DN1s and DN3s were the strongest sleep-promoting neurons. Not surprisingly, the E cells were the strongest wake-promoting neurons (Figure 1c). Moreover, it was gratifying that all of the methods, including *trans-*Tango, indicate that E-cells are bona fide post-synaptic targets of the sleep-promoting APDNs, consistent with previous work (Guo et al., 2016). Interestingly, the same approach failed to identify M cells as a post-synaptic target as this previous work also indicated. This could indicate that the *in-vitro* inhibition of M cells by DN1s is via neuropeptide signaling, but this previous work pointed to DN1-derived glutamate for M cell as well as E cell inhibition (Guo et al., 2016). This suggests that this previously observed *in-vitro* inhibition of M cells by DN1 firing is indirect and that the DN1 inhibition of locomotor activity occurs principally through E cell inhibition.

In addition to the well-described activity-promoting E cells, we were also able to identify another wake promoting circadian neuron subset, PI-projecting DN1s that target to the PI region. It is worth noting that the R18H11 driver previously only identified the sleep-promoting DN1s and did not identify the wake-promoting DN1 subset (Guo et al., 2016). Yet we show here that *Clk856-GAL4 ∩ R18H11-LexA* labels both activity- and sleep-promoting DN1s (Figure 2a and c). The discrepancy can be explained if the R18H11 driver expresses too weakly in the CRY-negative DN1s to be visualized by GFP staining, i.e., there is very little PDFR expression in these neurons (Im et al., 2011). However, that LexA driver is used here to express FLP, which just needs remove the stop codon to ensure strong expression of CsChrimson by *Clk856-GAL4* even within CRY negative DN1s.

The heterogeneity of DN1s with two different projection patterns and behavioral outputs may parallel the expression patterns of key molecules like CRY and PDFR. The CRY+ PDFR+ sleep-promoting DN1s are poised to integrate sensory input from the environment to regulate arousal and sleep/locomotion (Yadlapalli et al., 2018). This is not only because they express the photoreceptor CRY but also because they express PDFR and have a posterior projection to communicate with the core pacemaker M and E cells. In addition, a very recent paper reported that the sleep-promoting DN1s closely monitor environmental temperature to adjust the fly locomotor/sleep pattern (Yadlapalli et al., 2018).

CRY and PDFR are always co-expressed in circadian neurons (Im et al., 2011), so CRY-negative DN1s should also be PDFR-negative. This explains “why” these PI-projecting DN1s lack a posterior branch and fail to communicate with the dorsal projection of PDF+ sLNvs. Previous studies have reported that these PI-projecting CRY-negative DN1s connect to DH44+ PI neurons to regulate locomotor activity (Cavanaugh et al., 2014; Kunst et al., 2014). As wake-promoting E cells also direct target the PI region (Figure 1b) (Schubert et al., 2018), the PI neurons may receive wake signals from two different circadian sources to promote different activities, e.g., mating, feeding, foraging etc.

The sleep-promoting DN1s not only send posterior projections to the s-LNvs, but they also send anterior-projecting fibers to the central brain, hence the term APDNs. The anatomical data from trans-synaptic labelling and expansion microscopy indicate that the APDNs connect to EB ring neurons via the TuBus. Interestingly, functional calcium imaging shows that the APDNs can evoke dramatic calcium oscillation in specific EB Ring neurons via TuBu_sup_.

Further optogenetic manipulations demonstrate that the APDN-TuBu_sup_-EB circuit modulates arousal levels, consistent with previous results showing that these ring neurons enhance arousal threshold (Lebestky et al., 2009). Another study identified those DN1s as a “STOP”-promoter, because their acute activation with the same *R18H11-GAL4* driver induced quiescence in a group of flies, indicating DN1 activation may also act though the EB to increase arousal threshold for social interactions (Robie et al., 2017). These anatomical and functional data show that this singular circuit can cause the broad sweep of changes associated with the transition from sleep to wake (Figure 8).

**Figure 8.**
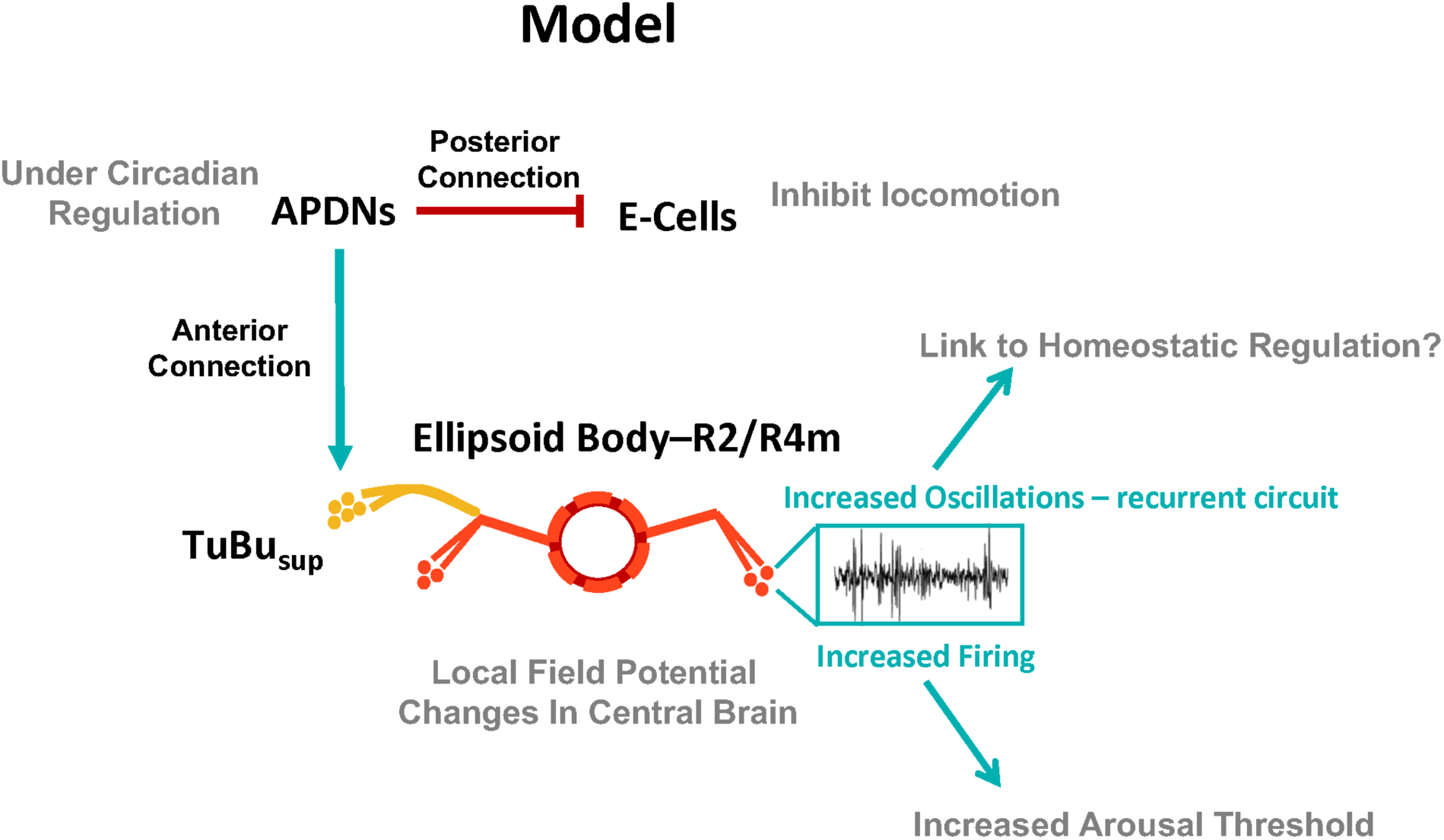
Model for the role of APDN-TuBu_sup_-EB-Ring neurons in sleep-wake arousal regulation. APDNs regulate sleep by two pathways. They decrease activity by inhibiting E cells through posterior projection. They also increase sleep-wake arousal threshold by activating TuBu_sup_ neurons and give rise to oscillations in EB-R2/R4m. The increased oscillation may reflect the recurrent circuit of EB and result in continuous increased arousal threshold during sleep as well as homeostatic sleep rebound.

EB neurons are essential for a variety of functions. They include integrating visual sensory input (Omoto et al., 2017), modulating attention and navigation (Sun et al., 2017) as well as regulating locomotor activity, arousal threshold (Lebestky et al., 2009) and sleep homeostasis (Liu et al., 2016). Another possible function for the conserved APDN-TuBu_sup_-EB circuit is that the APDNs provide circadian information to help invertebrates process sensory cues. For example, during monarch butterfly (*Danaus plexippus*) migration, circadian neurons can calibrate the time-compensated sun compass orientation, which contributes to navigation (Froy et al., 2003). Other diurnal insects like flies are able to switch behaviors based on the position of a bright celestial body like the sun. Interestingly, DN1s fire more during the morning comparing to the evening, so the CRY+ APDNs may provide photoperiod as well as time of day information to the central brain. Moreover, during nighttime migration or other nighttime activities when there is minimal ambient light (like during a new moon), the APDNs via the TuBu may be the major temporal input to the EB.

The induction of potent EB Ring neuron calcium oscillations by APDN activation is intriguing (Figure 6d-f). Although there is nothing known about the neuronal mechanism underlying these oscillations nor their purpose (if any), recent work in flies reported increased 7–10 Hz oscillations within the central brain during spontaneous sleep. Although this is much more rapid than mammalian delta waves (Yap et al., 2017), it is tempting to connect these oscillations as well as those we have observed to mammalian sleep. Moreover, it will be interesting to test whether APDN firing can produce similar oscillations or local field potential changes in other sleep-relevant central brain regions like dFSB. We speculate that these calcium oscillations reflect an organized recurrent circuit of the central complex (Donlea et al., 2018), which keeps EB ring neurons in a firing mode that influences their previously described effects on arousal threshold and sleep homeostatic regulation (Figure 8) (Liu et al., 2016).

Finally, we note that the EB may participate in modulating arousal threshold not just for shortening sleep latency or enhancing sleep maintenance. Increasing the central brain input threshold may help filter out unrelated external sensory input or unwanted stimuli to improve attention and ultimately animal survival. Given the role of the EB in vision-centric behaviors and navigation, it should be possible to test whether APDN input can improve fly attention in a visual discrimination task.

## EXPERIMENTAL PROCEDURES

### Fly strains

All flies were reared on standard cornmeal/agar medium supplemented with yeast. The adult flies were entrained in 12:12 light-dark (LD) cycles at 25°C. The fly lines used in the study were listed below: *UAS-GtACR1* was from Dr. Adam Claridge-Chang; *Clk4.1M-GAL4* was from Dr. Paul Hardin; *trans-* Tango related flies was from Dr. Gilad Barnea; *UAS-TNT* were from Dr. Hubert Amrein; *UAS-FRT-STOP-FRT-CsChrimson.mVenus* were from Dr. Bruce Baker; *CRY-GAL80* are described by Stoleru et al; *LexAop-P2X2*, *UAS-P2X2* and *Clk856-GAL4* were from Dr. Orie Shafer. The following lines were ordered from the Bloomington Stock Center: *R18H11-GAL4 (48832), R18H11-LexA (52535),UAS-CsChrimson.mVenus (55136), UAS-Denmark (33064), UAS-GCaMP6f (42747), QUAS-GCaMP3 (52231), UAS-syt-GFP (33064), R19C05-LexA (52541), R43D05-LexA (54147), R18H11-LexA (52535), R73B06-LexA (54228), R64A11-LexA (52825), R81E08-LexA (54377), R55C09-LexA (53510), R19A11-LexA (54772), R16C05-LexA (54457), R70E02-LexA (54227), R70G01-LexA (52855), R66B05-LexA (54917), R77H08-LexA (54392), R12G04-LexA (52448), R67F03-LexA (61585), R65D05-LexA (53625), R67F03-GAL4 (39448), R76B06-GAL4 (48327), R76B06-LexA (54225), R78B07-GAL4 (39989), LexAop2-FLP (55820), LexAop2-IVS-CsChrimson.mVenus (55138), LexAop2-IVS-NES-jRCaMP1b-p10 (64428), R41A08-GAL4 (50108), R10B11-GAL4 (48247), VT038828-GAL4 (v201975)* was from Vienna Drosophila Resource Center.

### Locomotor activity and statistical analyses

Locomotor activity of individual male flies (aged 3-7 days) was measured with video recording Flybox under 12:12 LD conditions. The analysis for activity and sleep was performed with pysolo and a signal-processing toolbox implemented in MATLAB (MathWorks, Natick, MA).

All statistical analyses were conducted using IBM SPSS software. The Wilks-Shapiro test was used to determine normality of data. Normally distributed data were analyzed with 2-tailed, unpaired Student’s t-tests, one way analysis of variance (ANOVA) followed by a Tukey-Kramer HSD Test as the post hoc test or two-way analysis of variance (ANOVA) with post-hoc Bonferroni multiple comparisons. Data were presented as mean behavioral responses, and error bars represent the standard error of the mean (SEM). Differences between groups were considered significant if the probability of error was less than 0.05 (P < 0.05).

### Arousal thresholds assay

For mechanical stimulation, individual flies from different groups were loaded into 96-well plates and placed close to a small push-pull solenoid equipped inside Flybox. The tap frequency of the solenoid was directly driven by an Arduino UNO board (Smart Projects, Italy). 10 taps (1 Hz) with 30 min interval was used as a strong stimulus. Arousal threshold was measured during 24 hr in day 4. The movement of flies before and after the stimulus was monitored by the web camera and the recording pictures with 10s interval were processed by MATLAB to calculate the locomotion curve of aroused flies.

### Feeding of retinal

All trans-retinal powder (Sigma) was dissolved in EtOH as a 100 mM stock solution. 100 µl of this stock solution was diluted in 25 ml of 5% sucrose and 1% agar medium to prepare 400 μM of all trans-retinal (ATR) food. For CsChrimson and GtACR1 experiments, flies were transferred to 96-well plates loaded with 300μl food (5% sucrose and 2% agar) containing 400 μM all trans-retinal (Sigma) for at least 3 days prior to any optogenetic experiments.

### Optogenetics and video recording system

The behavioral setup for the optogenetics and video recording system is described in prev ious publication. Flies were loaded into white 96-well Microfluor 2 plates (Fisher) containing 5% sucrose and 1% agar food with 400 μM ATR. Back lighting for night vision was supplied by an 850 nm LED board (LUXEON) located under the plate. Two sets of high power red LEDs (630 nm) and green LEDs (540 nm) mounted on heat sinks were symmetrically placed above the plate to provide light stimulation.

The angle and height of the LEDs were adjusted to ensure uniform illumination. The voltage and frequency of red light pulses were controlled by an Arduino UNO board (Smart Projects, Italy). We used 627 nm red light pulses at 10Hz (0.08mW/mm^2^) to irradiate flies expressing the red-shifted channelrhodopsin CsChrimson. Fly behavior was recorded by a web camera (Logistic C910) without an IR filter. We used time-lapse software to capture snapshots at 10 second intervals. The LD cycle and temperature was controlled by the incubator, and the light intensity was maintained in a region that allowed entrainment of flies without activating CsChrimson or GtACR1. Fly movement was calculated by Pysolo software and transformed into a MATLAB readable file. 5 pixels per second (50% of the Full Body Length) was defined as a minimum movement threshold. The activity and sleep analyses were performed with a signal-processing toolbox implemented in MATLAB (MathWorks) as described above.

### Fly brain immunocytochemistry and Pro-ExM

Flies were fixed in PBS with 4% paraformaldehyde and 0.008% Triton X-100 for 3hr at room temperature. Fixed flies were then washed in PBS with 0.5% Triton X-100 and dissected in AHL. The brains were blocked in 10% goat serum (Jackson Immunoresearch, West Grove, PA) and subsequently incubated with primary antibodies at room temperature for 36hr. A rabbit anti-DsRed (Clonetech, 1:200), a mouse anti-PDF (DSHB, 1:1000), a mouse anti-HA (Covance, MMS-101P; 1:250) and a chicken anti-GFP antibody (Abcam, 1:1000) antibody were used as primary antibodies for experiment purpose. After washing with 0.5% PBST three times, the brains were incubated with Alexa Fluor 633 conjugated anti-rabbit and Alexa Fluor 488 conjugated anti-chicken (Molecular Probes, Carlsbad, CA) at 1:500 dilution for three hours at room temperature. The brains were washed with 0.5% PBST three times before proceeding with either the traditional mounting method or the Pro-ExM protocol. For regular immunohistochemistry, the brains were mounted in Vectashield Mounting Medium (Vector Laboratories, Burlingame, CA) and viewed sequentially in 1 μm sections on a Zeiss LSM 880 confocal microscope with a multi-immersion 25x objective.

For ExM-treated brains, the Pro-ExM protocol described in Tillberg et al., (2016) was conducted with the following modifications: The anchoring treatment comprised of the Acryloyl X – SE (Life technologies A20770; 1:100) resuspended in PBS solution for 24 hours. The brains were washed in PBS three times before the gelation. The Gelling Solution was added and incubated on ice for 45 minutes before pipetting the brains onto the Gel Chamber composed of a glass slide, spacers, and a coverglass. The assembled Gel Chamber with the brains were incubated at 37°C for about 2 hours. When the gel solidified, the embedded brains were trimmed away from the Gel Chamber and submerged in the Digestion Buffer with fresh Proteinase K (1:100) for 24 hours. Lastly, the brains were washed with excess ddH2O for at least three times, 20 minutes each. The embedded brains will then swell about 4 times the original size and will be nearly transparent. The expanded brains were placed on a glass bottom culture dish (MatTek Corp, P35GC-0-14-C) with dH2O. The brains were viewed sequentially in 1 μm sections on a Zeiss LSM 880 confocal microscope with a multi-immersion 25x objective. All the images were viewed post-hoc on ImageJ, including the z-depth color coding.

### Functional fluorescence imaging

For imaging experiments, adult fly brains were dissected in hemolymph-like saline (AHL) containing 108 mM NaCl, 5 mM KCl, 2 mM CaCl2, 8.2 mM MgCl_2_, 4 mM NaHCO_3_, 1 mM NaH2PO_4_-H_2_O, 5 mM trehalose, 10 mM sucrose, 5 mM HEPES; pH 7.5. The dissected brain was then pinned on to a layer of Sylgard (Dow Corning, Midland, MI) silicone under a 1ml bath of AHL contained within a recording/perfusion chamber (Warner Instruments, Hamden, CT) and bathed with room temperature AHL. Perfusion flow was established over the brain with a gravity-fed ValveLink perfusion system (Automate Scientific, Berkeley, CA). 2.5 mM ATP was delivered by switching the perfusion flow from the main AHL line to another channel containing diluted compound after 30 s-1min of baseline recording for the desired durations followed by a return to AHL flow.

Spinning Disk Confocal (Intelligent Imaging Innovations, Inc. Denver, CO) was used for GCaMP and R-CaMP recordings, Fluorescence signals were recorded at 2 Hz. ROIs were analyzed using commercial software SlideBook 6. The fluorescence change was calculated by using the formula: Max ΔF/F = Max (Ft – F0)/F0 × 100% or Ft/F0, where Ft is the fluorescence at time point n, and F0 is the fluorescence at time 0.

## AUTHOR CONTRIBUTIONS

F.G., M.H. and M.R. conceived and designed the experiments. F.G. and M.H. performed the behavioral experiments, immunostaining and functional imaging experiments. M.D., M.H. and F.G. performed the ExM experiments. F.G. and M.H. analyzed the data. F.G., M.H., M.D. and M.R. prepared the figures and wrote the paper.

## ACKNOWLEDGMENTS

We thank Katharine C. Abruzzi, Matthias Schlichting and Shlesha Richhariya for comments on the manuscript, and also thank Gilad Barnea, Bruce Baker, Adam Claridge-Chang and the Bloomington Stock Center for reagents. We are grateful to the members of the Rosbash Lab and our neighbors in the Griffith Lab for generous help, advice and discussions.

